# Relatedness of *Gardnerella* prophages and their endolysins

**DOI:** 10.64898/2026.09.28.754664

**Authors:** Brandon S. Maust, Kira A. Griswold, Monique Khim, Sandhya Subramanian, Bart Staker, Peter J. Myler, Brian R. Kullin, Anna-Ursula Happel, Arvind Varsani, Heather B. Jaspan

## Abstract

The *Gardnerella* genus includes a complex sub-taxonomy of bacterial species and subgroups which have been implicated in bacterial vaginosis. We analyzed *G. leopoldii, G. swidsinski, G. piotii*, and *G. vaginalis* genomes available in public databases and predicted proviral sequences using Cenote-Taker3. We clustered 266 prophage sequences into 140 viral genera level groupings that encode four distinct gene clusters and 19 different lysin proteins. Analysis of the bacterial and prophage sequence phylogenies reveals that while there were no specific prophage sequence lineages that infected each bacterial host, there is clustering of the prophage sequences by bacterial species within the *Gardnerella* genus.

**Impact Statement:** We found that many bacteriophages infecting different *Gardnerella* species are closely related, and that despite the diversity at the genome scale a small number of endolysin proteins were shared among a large number of prophages.

**Data Summary:** The bacterial genomes analyzed during the current study are available in the NCBI GenBank and GTDB repositories under the accession numbers listed in Supporting Data Table 1. Additional code for the analysis is available at https://github.com/bmaust/gardnerella_phage.

## Introduction

Bacterial vaginosis (BV) is the most common cause of vaginal dysbiosis among women of reproductive age and is associated with a range of adverse reproductive and sexual health outcomes.^1^ Although BV is a polymicrobial condition, species within the *Gardnerella* genus are considered key initiators of the biofilm associated with BV.^2,3^ The substantial heterogenicity within the *Gardnerella* genus, with multiple species and sub-species having distinct genomic content and associated virulence, has resulted in the recent formal and proposed taxonomic revisions of the genus.^4–7^ The biological differences among *Gardnerella* species and strains are clinically important but remain poorly understood.

Despite the central role of *Gardnerella* in BV, associated bacteriophages have been minimally characterized. Prophages are recognized as important drivers of bacterial evolution through horizontal gene transfer, host adaption, modulation of virulence, and shaping bacterial community structure.^8–11^ Previous work identified a high proportion of predicted prophages in *Gardnerella vaginalis* genomes from urinary isolates,^12^ suggesting frequent infection. A recent report of the bacterial pangenome within the genus reported two prevalent clusters of phage, but did not additionally characterize them^7^, further suggesting that prophages are common among *Gardnerella.* An analysis of CRISPR spacers in *Gardnerella* genomes found archived sequence fragments with homology to (presumably prophage in) genomes from other *Gardnerella* species and more distantly related bacteria,^13^ further evidence of *Gardnerella* phages’ ability to infect across strain or species boundaries.

Despite this evidence of widespread lysogeny, characterization of *Gardnerella* phage has been limited. To date, only one phage infecting *Gardnerella vaginalis* has been isolated and genomically characterized,^14^ and we could find no other additional published reports of successful prophage induction from *Gardnerella* cultures. The lack of cultured *Gardnerella* phages despite the prevalence of prophage-like sequences in genomes suggests that the diversity of *Gardnerella* phages is still underexplored.

Characterizing *Gardnerella* phages is of particular interest, as current standard of care antibiotic regimens for BV are limited by suboptimal response rate and increasing antimicrobial resistance.^1^ Further, recurrence is high, partly because *Gardnerella* biofilms reduce antimicrobial penetration and facilitate recolonization.^15,16^ Therefore, there is growing interest in novel therapeutics that selectively target *Gardnerella*.

Phage-derived recombinant endolysins might be a promising approach. Several *Gardnerella* endolysins have been recently characterized, with the goal to be developed as a potential treatment for BV.^17^ For example, drug candidate PM477 disrupts *Gardnerella* biofilms while not affecting beneficial vaginal bacteria.^17^

Here, we analyze genomes from multiple *Gardnerella* species to systematically identify prophages and encoded lysin proteins. By providing a comprehensive overview of bacteriophage diversity within the genus, we aim to examine evidence for ongoing bacteriophage evolution rather than fossilized carriage within the bacterial genomes.

With potential application for therapeutics, we also examine the degree to which the phage genomes show specialization by bacterial host lineage.

## Materials and Methods

### Phage prediction and annotation

We collected genome sequences for vaginal *Gardnerella* species (*G. piotii*, *G. leopoldii*, *G. swidsinskii*, *G. vaginalis*) from GTDB^18^ and NCBI GenBank^19^ (Supporting Data 1). For sequences with conflicting taxonomic designations, we chose the GTDB entry over NCBI or newer proposed taxonomy (i.e. in Bouzek et al.^7^). We confirmed there were no duplicate bacterial sequences using FastANI v1.34.^20^ Cenote-Taker^21^ v3.4.1 was used to identify putative prophage sequences and assign taxonomy, using the options --prune_prophage T and --lin_minimum_hallmark_genes 2. Viral contigs were manually inspected for bacterial genes to confirm Cenote’s pruning.

We compared the phage genomes within and between *Gardnerella* species using CD-HIT^22^ at 99% average nucleotide identity (ANI) and 80% length difference cutoff to determine duplicates. We used CheckV^23^ v 1.0.3 to estimate the genomic completeness of the sequences and retained only those of medium or higher quality (≥ 50% complete). These phage sequences were clustered at a 70% average nucleotide identity threshold with CD-HIT to define genus-level clusters.^24^ We also compared the distribution of phage ANI distances within each *Gardnerella* species using PERMANOVA as implemented in the adonis2 function of vegan v2.7.3. Phage sequence genome maps were created with gggenomes^25^ v1.1.3 using Phrog^26^ categories.

Given that the PhiX-174 bacteriophage sequencing control used in Illumina sequencing frequently contaminates metagenome assembled genomes,^27^ we used NCBI blastn^28^ v 2.17.0+ to measure homology between the identified prophage sequences and its reference genome (NCBI RefSeq NC_001422).

### Protein annotation and analysis

Coding sequences (CDS) were annotated in all 396 of the predicted prophage sequences with predicted function by Pharokka^29^ v 1.8 using the PHROG,^26^ VFDB,^30^ and CARD^31^ databases via MMseqs2^32^ and PyHMMER.^33^ We used Phold^34^ v 1.2.5 for additional functional prediction via structural homology using Foldseek,^35^ ProstT5,^36^ and Colabfold.^37^ We used anvi’o^38^ v9 to cluster CDS and visualize phage pangenomes using –minbit 0.5 and –mcl-inflation 10.

To analyze lysins, we extracted all predicted CDS with PHROG category of “lysis” and not the function “holin” with confidence of medium or high. These putative lysin CDS were compared within and between *Gardnerella* species using CD-HIT^22^ at 99% amino acid identity to detect duplicates. We also used a threshold of 85% amino acid identity to cluster related lysins for downstream analysis. Structures were predicted for the protein sequences with Boltz2^39^ v 2.2.0 and Pfam^40^ functional domains were annotated using InterPro^41^ v 109. Structures were visualized using UCSF Chimera X^42^ v 1.12.

### Phylogenetic inference

We generated a proteomic phylogeny of the predicted phage sequences using VIPTree.^43^ For the bacterial sequences we used the *cpnDB*^44^ v 13.0 with NCBI blastn to extract the corresponding region from each *Gardnerella* genome, then used clustal^45^ to align the nucleotides and PhyML^46^ with the generalized time reversible nucleotide substitution model to reconstruct a phylogeny. The two trees were co-visualized using phytools.^47^ We computed the mean pairwise distance (MPD) and mean nearest taxon distance (MNTD) on the viral tree using the ses.mpd and ses.mntd functions of picante v 1.8.4, both with 999 runs permuting the taxa labels to generate the null.

### Accumulation curves

We calculated accumulation curves using vegan^48^ v2.7-3 for predicted viral genomes at 95% ANI (the species threshold^24^), and 85% amino acid identity for the annotated lysin and major tail protein CDS.

## Results

### Gardnerella frequently hosts predicted prophage

We identified 297 putative prophage sequences in 198 of the 454 *Gardnerella* genome sequences available in public databases (**Table 1**). Among bacterial genomes with any virus-like elements identified, the median number identified was 1 with a maximum of 8. Four alphapapillomaviruses genomes were identified and removed from further analysis as likely assembly errors. We found no evidence of PhiX-174 sequence contamination in the identified virus sequences.

The average length of the putative prophage contigs was 37 kilobases (range: 1 kb-99 kb) with a mean of 43 open reading frames (ORFs) (**Figure 1**). A substantial majority of the genomes (95%) were predicted to belong to the class *Caudoviricetes*, with an additional 3 (1%) in the class *Megaviricetes*, and 11 (3.6%) that were unable to be taxonomically classified.

**Figure 1:**
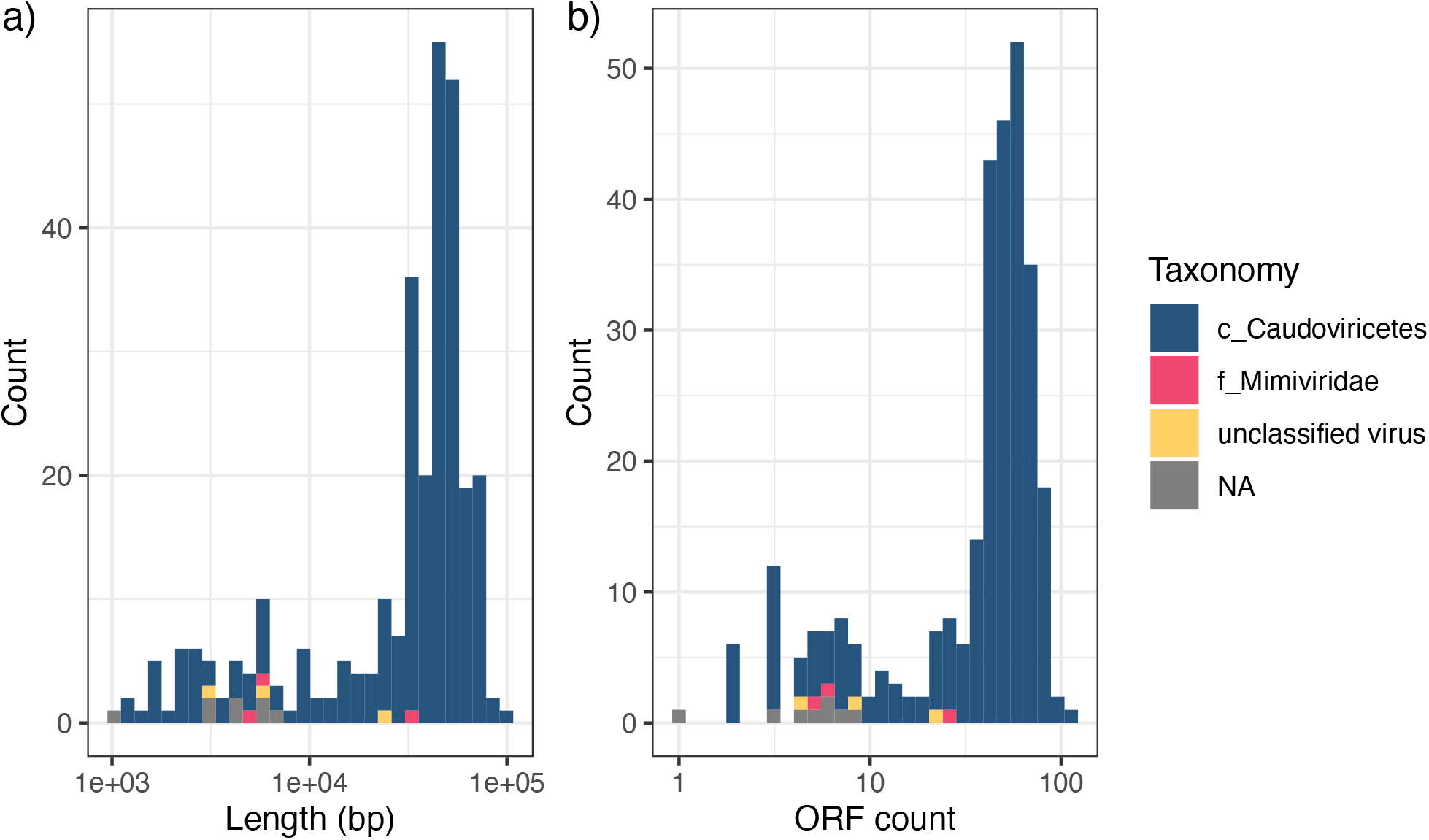
a) Length of the virus-like contigs as a histogram, colored by taxonomic assignment. b) Count of ORFs per contig as a histogram, colored by taxonomic assignment

Of the predicted prophage sequences, 31 shared 99% average nucleic acid identity (ANI) with at least one other sequence, and thus 266 unique phage sequences were identified. All the clusters of highly similar sequences were within different host bacterial sequences within the same *Gardnerella* species; none of them were duplications within the same bacterial genome.

We removed sequences labeled by CheckV^23^ as low-quality or not-determined, leaving 213 sequences designated as medium quality (n=56), high quality (n=156), or complete (n=1) for downstream analysis. Further clustering the predicted prophage sequences at 70% ANI resulted in 140 representative genus-level sequences.

### Gardnerella prophage sequences are loosely related

Clustering by both CDS co-occurrence and ANI segregated the sequences into four groups (**Figure 2**). There were two distinct clusters (**Figure 2**, upper right) with minimal overlap in gene clusters or nucleotide homology. The first cluster includes two highly similar sequences from bacterial genomes that were sequenced in the same project from the vaginal secretions of participants with BV in China.^4^ The second, largest, cluster includes the previously reported vB_Gva_AB1 and has members from each *Gardnerella* species (**Figure 2**). There were also two less distinct clusters with some shared CDS (lower left), including the recently-described pGamma^7^ (**Figure 2**).

**Figure 2:**
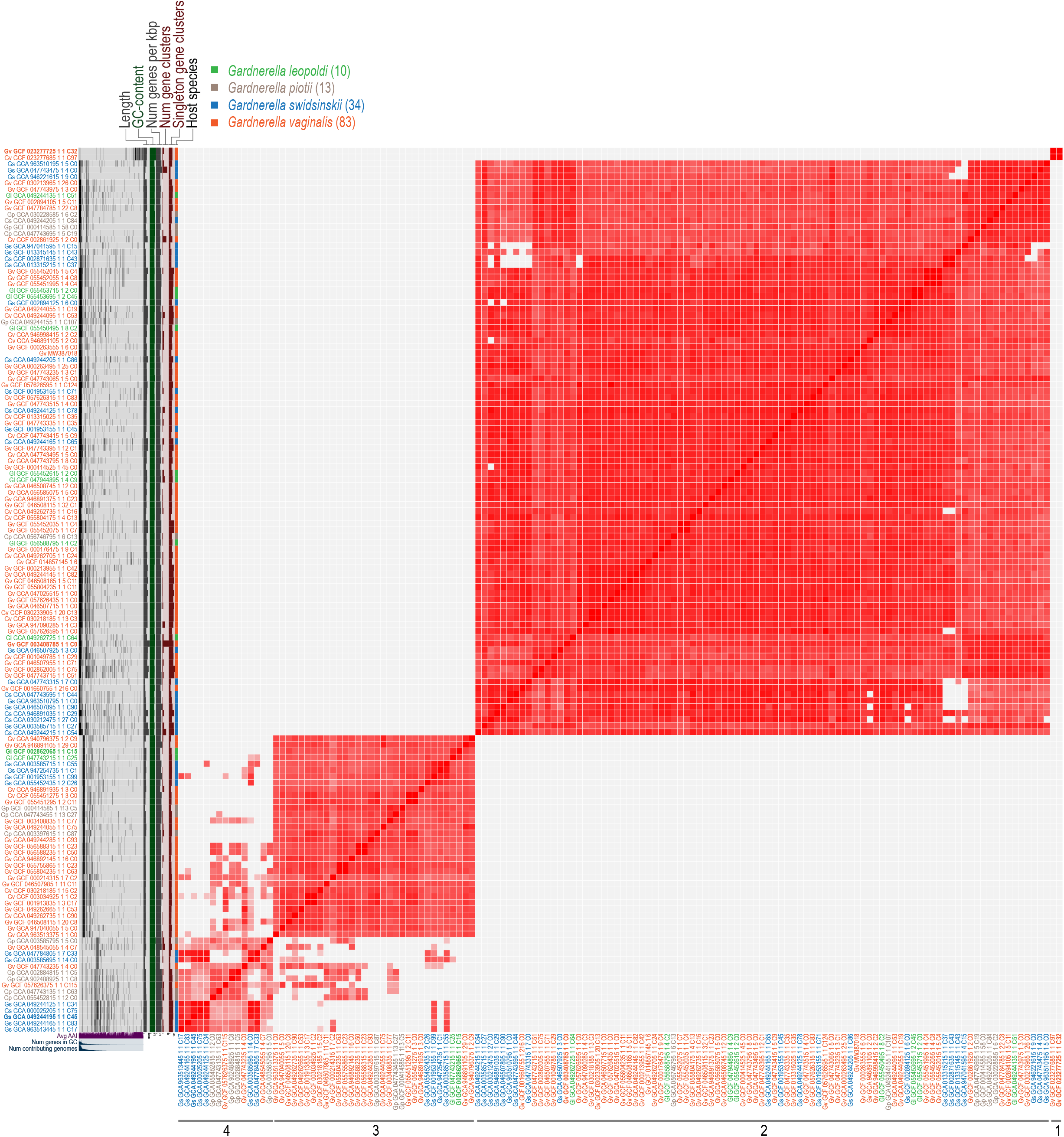
Gene clusters (left) with graph of gene cluster membership across the genome for each sequence next to its name followed by graphs of summary statistics (genome length, average GC content, genes per kilobase, and the total number of shared and singleton gene clusters) and bacterial host species. The representative sequences from each cluster used in Figure 3 are indicated in bold. To the right in red is the pairwise average nucleotide identity (FastANI) with the cluster memberships below. At the bottom are shown summaries of Amino Acid Identiy (AAI) across the genome, gene cluster content, and gene cluster sequence membership.

The low level of shared gene content between these bacteriophage genome groups supports their distinctiveness. The prevalence of gene clusters within the groups rarely met the >95% thresholds used to define core genomes in bacteria and fewer than 4% of gene clusters were present across even 50% of viral sequences (**Table 2**). Within the clusters defined above there was more consistency in gene content between contigs. In the larger sequence cluster (2), a single gene cluster comprising 92 tail protein genes and 5 unannotated genes was present in 95% of sequences (Supplemental Data 1). In the smaller sequence cluster 3, ten gene clusters were found at 100% prevalence and encoded proteins for replication and assembly with no additional gene clusters above 95% prevalence at which point additional structural and assembly proteins were added (Supplemental Data 1). In sequence cluster 4, one gene cluster of structural proteins was found at 100% prevalence and no additional gene clusters were found above 90% prevalence (Supplemental Data 1).

**Table 1:**
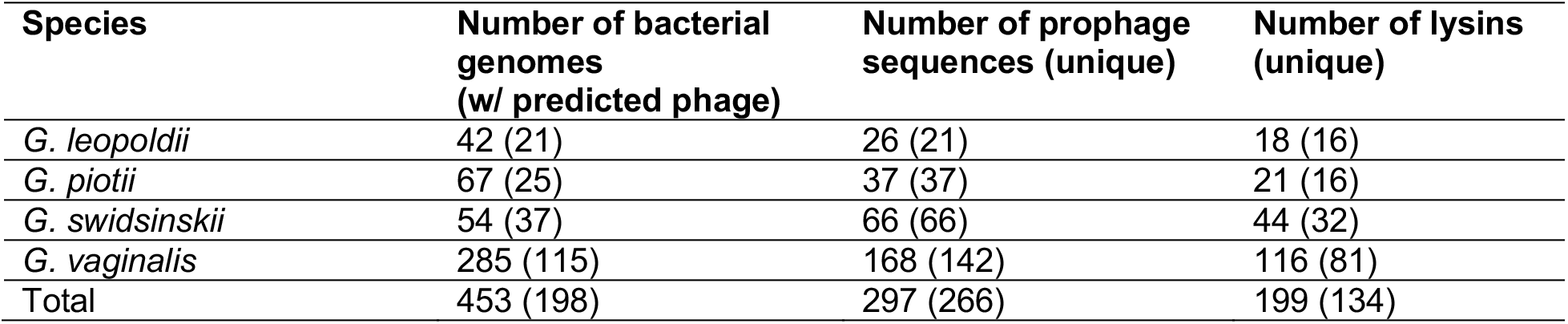
Counts of bacterial genomes from NCBI and GTDB, the raw counts of predicted prophage and the number of unique sequences (clustering the predicted prophage sequences at 99% ANI), and the number of annotated lysin CDS among the predicted prophage along with the number unique (clustering at 99% AAI).

**Table 2:** Gene cluster frequencies (% of non-singleton clusters identified in each group) by Anvi’o gene cluster group. Group 1 is excluded as it only contained 2 sequences.

| Prevalence (%) | All | Group 2 | Group 3 | Group 4 |
| --- | --- | --- | --- | --- |
| 100 | 0 | 0 | 6.2 | 0.5 |
| 99 | 0 | 0 | 6.2 | 0.5 |
| 95 | 0 | 0.2 | 14.3 | 0.5 |
| 90 | 0 | 3.8 | 18 | 0.5 |
| 75 | 0 | 6.2 | 18.6 | 3.9 |
| 50 | 3.7 | 7.6 | 29.2 | 19.7 |

Testing the ANI between the prophage sequences found in each bacterial species revealed no statistically significant differences (PERMANOVA p=0.06 R^2^=0.04). Visualizing the distances by Principle Coordinates Analysis (PCoA) reveals four obvious clusters of sequences corresponding to the high pairwise identity clusters identified in Figure 2 with broad distribution of the bacterial species through the clusters (**Figure 3A)**.

**Figure 3.**
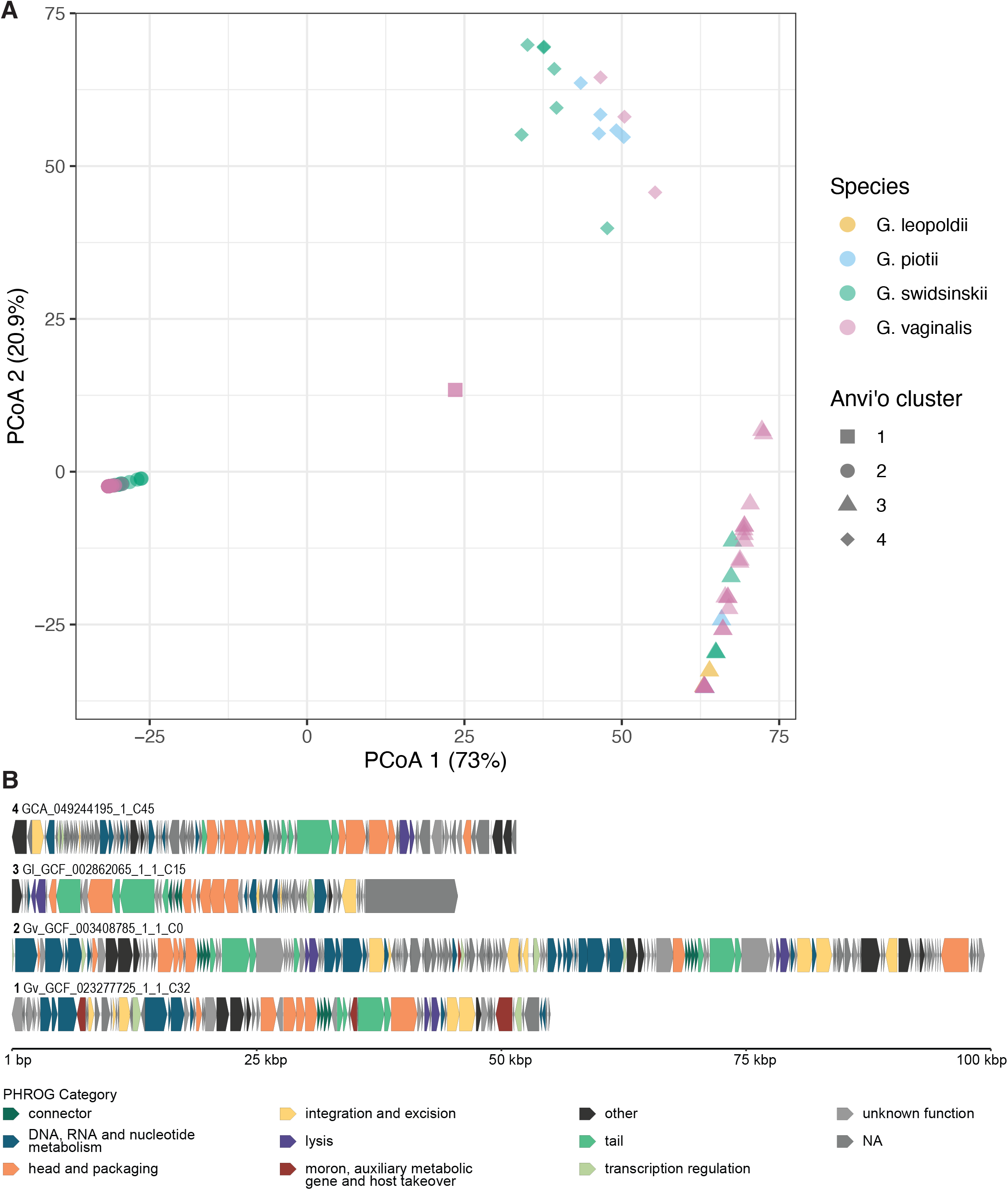
Phage genomes. **A)** PCoA of FastANI distances between the 140 prophage contigs. Host bacterial species is denoted by color and the cluster from Figure 2 is denoted by shape. **B)** Genome maps of a representative sequence from each cluster of *Gardnerella* phage in Figure 2. The cluster number is indicated in bold at the beginning of the sequence name. CDS are colored by PHROG category.

Examining a representative sequence (the longest) for each sequence cluster showed CDS with diverse PHROG categories, consistent with the prophages being functionally intact and unlikely to represent fragmented genomes from sequencing or assembly errors (**Figure 3B**). Aside from the small cluster of two viral sequences, prophages identified in multiple *Gardnerella* species were clustered together, suggesting that viral diversity is not constrained along bacterial host taxonomic boundaries.

### Evidence for independent evolution of bacterial sequences and putative prophages

To further examine the relationship between the host bacterial and prophage sequence phylogenies, we separately estimated phylogenies using the sequences from each kingdom and graphed them together (**Figure 4**). In the bacterial *cpn60* phylogeny, the bacterial species are monophyletic (with the exception of occasional *Gardnerella vaginalis* designated genomes intermixing with the more recently described species), but the virus-like sequences found in the bacterial genomes did not visually segregate in the viral phylogeny by their origin host.

**Figure 4:**
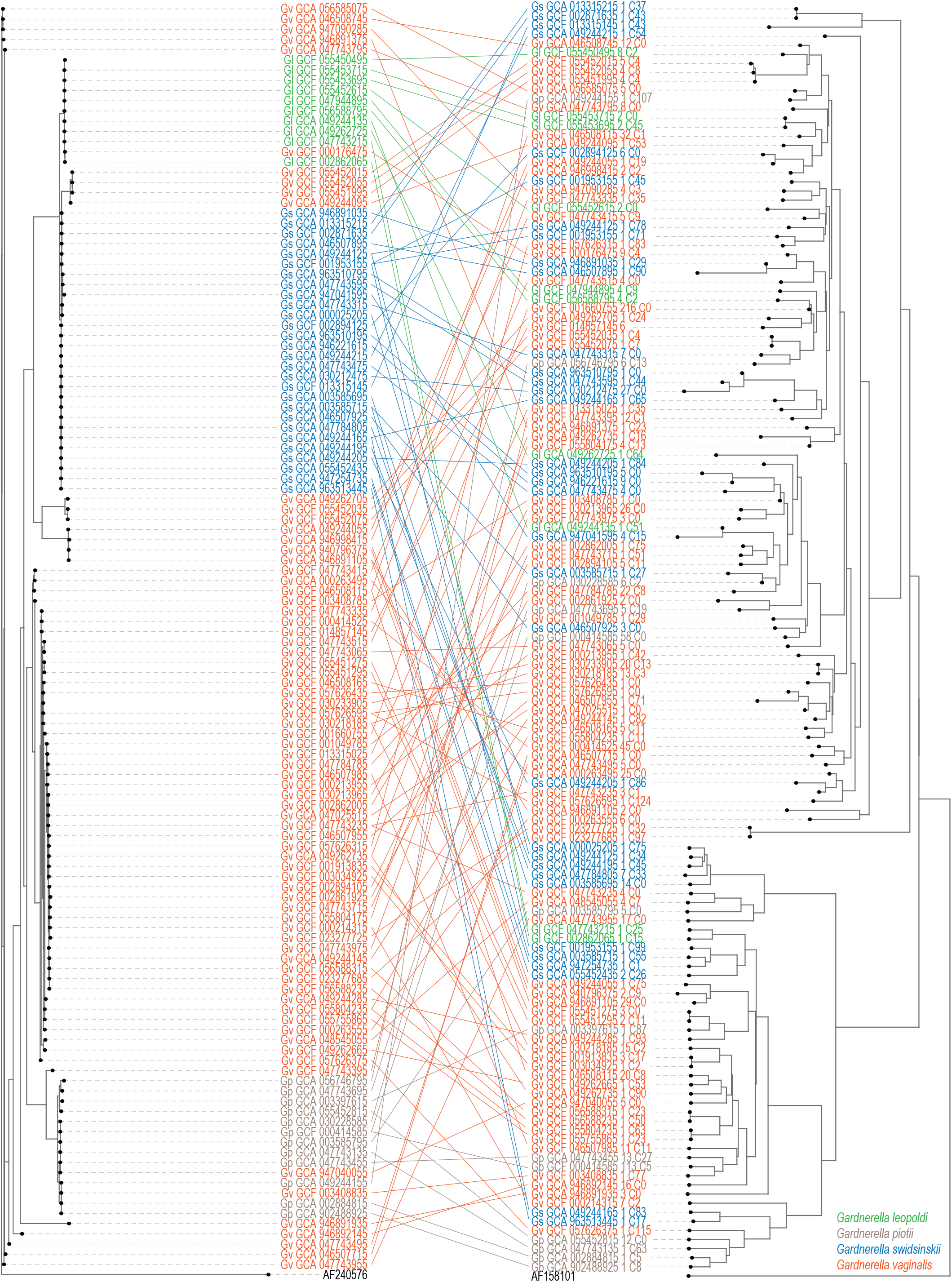
Phylogenetic relatedness of *Gardnerella* and prophage genomes. Bacteria (left, colored by and labeled with abbreviated with the first letter of the genus and species at the start of the RefSeq genome assembly ID) are connected to their predicted prophages (right, colored and labeled with the host RefSeq bacterial genome assembly ID and the contig, and virus-like sequence within the contig as designated by Cenote-taker). *Bifidobacterium longum* (GenBank accession AF240576) was used as an outgroup for the bacteria, and the *Escherichia coli* T4 phage (GenBank accession AF158101) was used for the phage.

To more rigorously evaluate whether bacterial host species were partitioned across the viral phylogeny, we examined the MPD and MNTD for each of the host species.

*Gardnerella vaginalis* prophages showed clustering in both MPD and MNTD and showed several distinct clades in the viral phylogeny with consistent labels (**Table 3** and **Figure 4**). *G. swidsinski* showed significant overdispersion of MPD and clustering of MNTD (**Table 3** and **Figure 4**), consistent with its phages appearing in multiple tightly related clusters. *G. leopoldii* and *G. piotii* did not show evidence of significant MPD clustering or dispersion, but both showed significant MNTD clustering (**Table 3** and **Figure 4**). Consistent with our other results, this suggests recent divergent evolution of these phages rather than a distant evolutionary segregating event as presumed for the bacteria. Although there were more *G. vaginalis* sequences in our data set, finding clustering at both levels is consistent with both the horizontal and vertical transfer characterizing phage inheritance.^49^

**Table 3:**
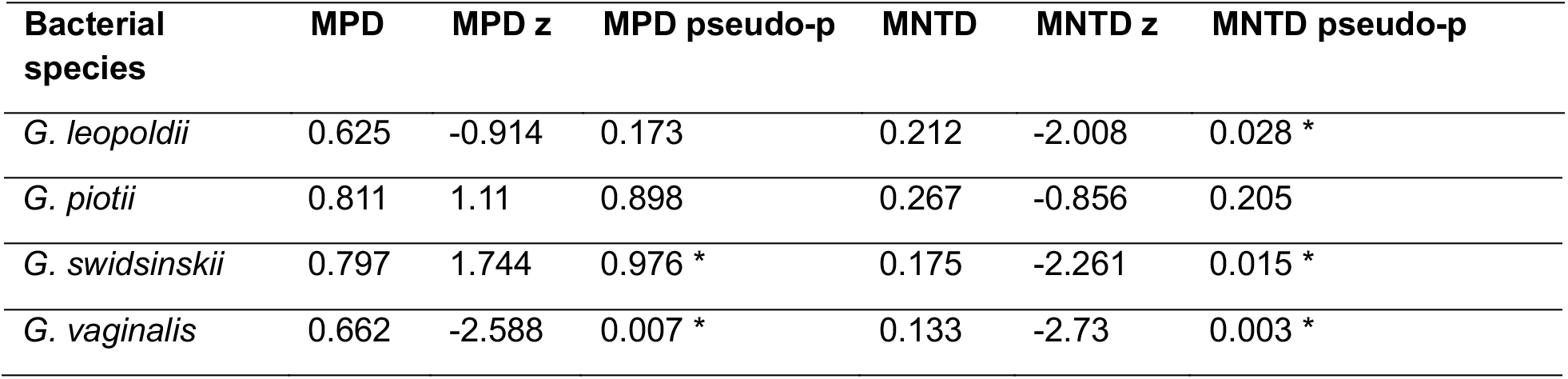
Mean pairwise distance (MPD) and mean nearest taxa distance (MNTD) measures for the viral sequences found in each *Gardnerella* species. Asterisk indicates statistical significance with one-tailed pseudo-p >0.95 or <0.05.

### Lysin proteins show high similarity across diverse predicted viruses

We focused additional attention on the endolysins predicted in the prophage sequences. The 134 unique endolysin proteins from 199 CDS in the predicted phage sequences clustered into 19 families, with the largest two clusters containing lysin sequences found in all four *Gardnerella* species. Two protein sequences were < 200 amino acids and had no domains identified by InterPro. Each of the 17 remaining sequences had a domain annotated with peptidoglycan hydrolytic activity, typically at the N terminal (**Figure 5**). In 12 (80%) of them this was a glycosyl hydrolase family 25 domain. In five sequences there was an annotation of a C-terminal domain for binding peptidoglycan.

**Figure 5:**
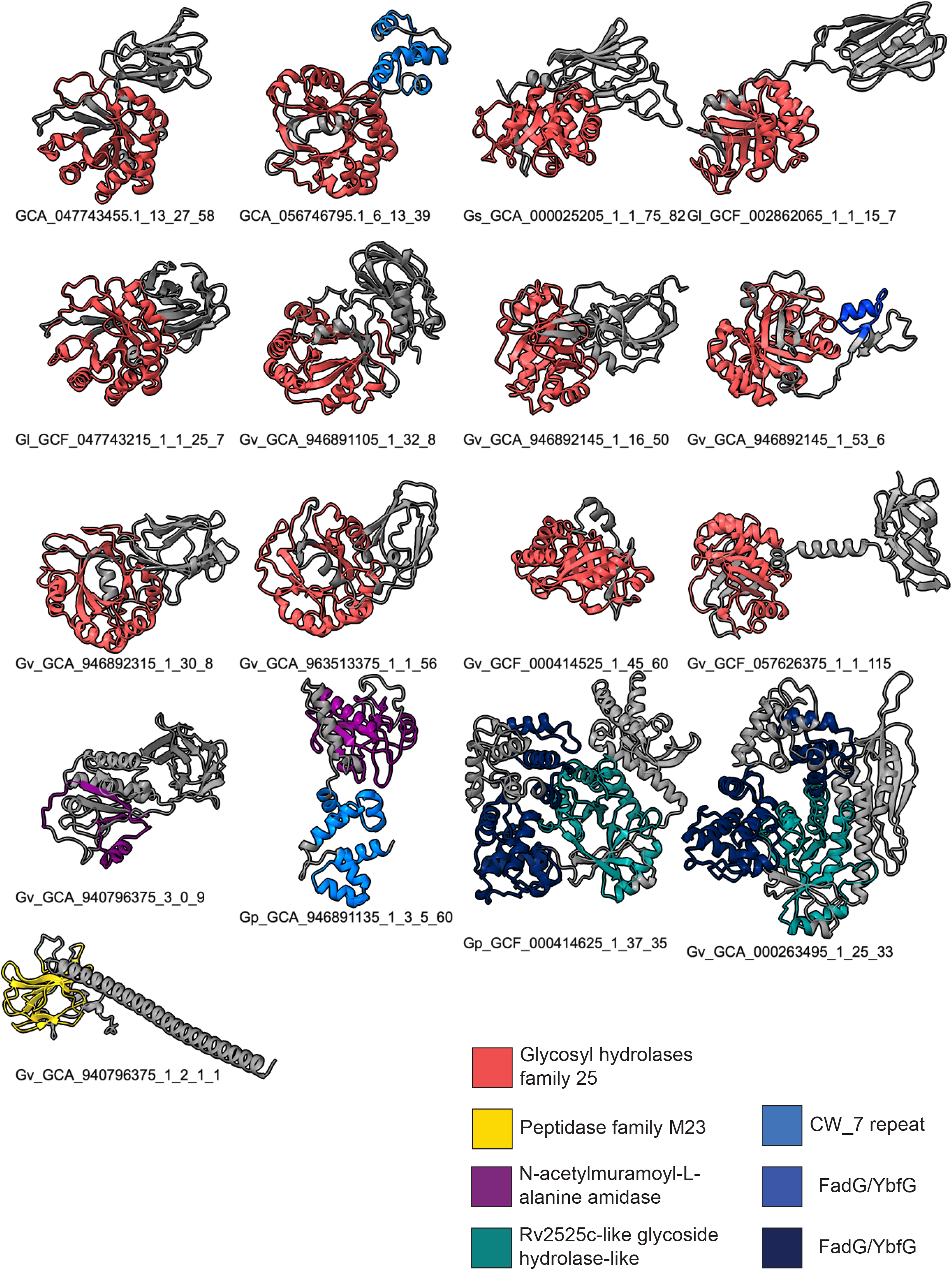
Endolysin predicted structures with Pfam domain annotations. Structures were predicted with Boltz2 and domains annotated by InterPro were added in ChimeraX. Binding domains are indicated in blue colors and catalytic domains in other hues.

### Prophage diversity but not lysin or structural protein diversity remains under-characterized

To estimate the representation of total prophage diversity from the sequences described here, we calculated an enrichment curve for viral genomes, lysin proteins, and major tail proteins **(Figure 6**). The viral genomes showed a continuously positive slope even including all sequences predicted from published bacterial genomes, while the two proteins showed saturation by as few as 25 genomes.

**Figure 6:**
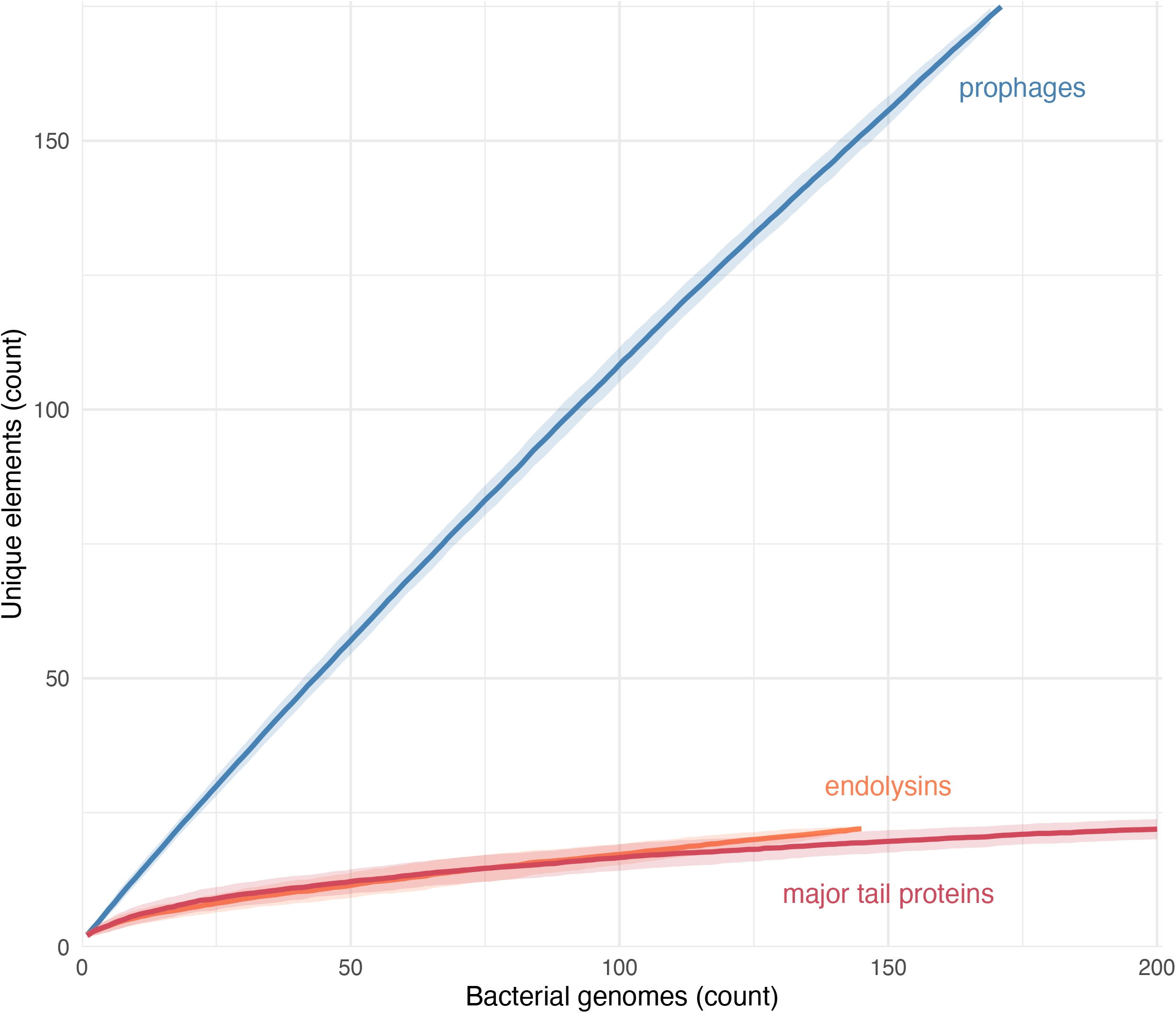
Enrichment curves for prophages, endolysins, and major tail proteins. Phage sequences (blue) were clustered at the species threshold of 95% ANI and the lysin (orange) and major tail (red) proteins at 85% AAI.

## Discussion

We used published *Gardnerella* genome sequences to identify predicted prophage sequences. As in a previous report focused on urinary isolates,^12^ we found putative prophage sequences in 44% of the *Gardnerella* genome sequences that we analyzed. Since not all available *Gardnerella* genome sequences are complete, this is likely an underestimate of the total prevalence of *Gardnerella* prophage. The lack of shared prophage sequences between *Gardnerella* species is consistent with phage host specificity at this level, either due to limitations on viral entry or superinfection exclusion through bacterial or viral mechanisms.

Most of the identified prophage sequences were classified in the *Caudoviricetes* class, which is consistent with the well-documented ability of such viruses to establish lysogeny. Among these sequences, we identified a large number of ORFs annotated as lysins, proteins that rupture the cell wall to allow progeny virions to be released during lytic replication. Despite the variation in nucleic acid sequence or these proteins, the shared catalytic domain annotations show relatively little variation, with the same Glycosyl hydrolase family 25 domain appearing in prophages from multiple *Gardnerella* host species. Finding fewer annotations for peptidoglycan binding or recognitions domains is likely a reflection of the limited repertoire of viral proteins in the annotation database.

The prophage proteome sequence phylogeny shows several closely related groups of viruses, somewhat more than previously reported^7,12^ - we think due to our inclusion of additional host bacteria and prophage sequences. There is no clear relationship between the bacteriophage and bacterial clades to suggest that host specificity shaped evolution at the level of the proteome. Phages from distant viral lineages were found in bacteria with closely related cpn60 chaperonin protein sequences, suggesting that the prophage sequences have been subject to independent evolutionary selection rather than carried along through bacterial evolution after a distant integration event, although it is also possible that other bacterial proteins would more concordantly segregate the bacterial phylogeny.

While the evolutionary role of prophages within *Gardnerella* genomes is unknown, their prevalence and diversity at both the level of nucleotide sequence and gene content argue against static carriage after an evolutionarily distant integration. The lack of a clear division of phage along the bacterial host species boundaries is likely a result of the complex evolutionary history between phages with both horizontal and vertical gene transfer. The recovery of closely related phage from bacteria across species boundaries is promising for medical applications of bacteriophage or their products, as it suggests that a small set of phages may be identified to infect a broad range of pathogenic *Gardnerella* strains.

Overall, these findings suggest that *Gardnerella* species have been infected by diverse lysogenic phages and that the current sequence databases have not sufficiently characterized this diversity. The relative conservation of lysin proteins suggests that one or a few may encompass the diversity across the breadth of observed *Gardnerella* phage, with positive implications for the feasibility of therapeutic application of phage proteins for this diverse bacterial genus. Additional work to characterize additional *Gardnerella* strains and their prophages remains important.

## Acknowledgements

The authors acknowledge Research Computing at Arizona State University and Research Scientific Computing at Seattle Children’s Research Institute for providing computing resources that have contributed to the research results reported within this paper.

## Supporting Data

Supporting Data Table 1: Accession numbers of all analyzed bacterial sequences

## Supplemental Data

Supplemental Data 1: Functional annotations for gene clusters

